# lncRNA DLEU2 acts as a miR-181a sponge regulated SEPP1 (may as a biomarker for sarcopenia) to inhibit skeletal muscle differentiation and regeneration

**DOI:** 10.1101/2020.04.15.025304

**Authors:** Yao Wang, Xue-ran Kang, Yisheng Chen

## Abstract

**Background:** Sarcopenia is a serious public health problem. The ceRNA network has been demonstrated vital in the development of skeletal muscle, but there is currently no effective method to assess the risk of sarcopenia. The purpose of this research is to create and authenticate a ceRNA pathway based on a predictive model of sarcopenia.

**Methods:** A clinical prediction model for sarcopenia was established using the RNAs (validated by clinical data) that are co-differentially expressed in the database, and a ceRNA network was constructed. The correlation analysis of each element in the ceRNA network was performed according to the clinical samples and the GTEX database, and the possible key ceRNA pathways were screened. C2C12 mouse myoblast Cells experiments were used to verify these ceRNA pathways.

**Findings:** Based on four molecular markers of SEPP1, SV2A, GOT1 and GFOD1, we developed a new model for predicting sarcopenia with well accuracy, and constructed a ceRNA network accordingly. Clinical sample showed that the expression levels of lncDLEU2, SEPP1, and miR-181a were closely related to the risk of sarcopenia. The C2C12 mouse myoblast cells were cultivated to verify that lncDLEU2 inhibits muscle proliferation and differentiation by acting as a miR-181a sponge regulated SEPP1.

**Interpretation:** Our research developed a highly accurate prediction tool for the risk of sarcopenia. These findings suggest that lncDLEU2-miR-181a-SEPP1 pathway inhibits muscle differentiation and regeneration. This pathway may uncover some new therapeutic targets for the treatment of sarcopenia caused by aging.

## Introduction

Sarcopenia has be closely associated with physical disabilities as well as the risk of diabetes and fractures characterized by a generalized and gradual loss of the function and strength of skeletal muscles. [1-7] With the increase of age, the incidence of sarcopenia has gradually increased and has become an important factor affecting the physical health of the elderly.[8-10] Skeletal muscle differentiation is affected by multiple signaling pathways. Myogenic regulation factor (MyoD, MyoG) is the core component of myogenic pathway.[11] With the development of sequencing, the role of lncRNA as a microRNA sponge to regulate miRNA’s ceRNA network in biological processes is becoming more and more widely recognized.[12-15] Several studies have demonstrated that long non-coding RNAs (lncRNAs), including MAR1, [16] H19,[17,18] MUMA,[19] Yam-1,[20] IRS1,[21] Malat1,[22] lncR-125b (TCONS_00006810),[23] and lnc-mg,[24] are associated with muscle differentiation and regeneration.

However, these previous researches mainly study the pathological role of the ceRNA network in sarcopenia, no quantifying method has been brought up to forecast the risk of sarcopenia; therefore, it is necessary based on several simple molecular markers to develop an reliable predictive tool and early interventions to reduce the risk of sarcopenia.

The purpose of this study was to create an effective tool for early predicting the risk of sarcopenia in patients based on several simple molecular markers. Based on this prediction model, we predicted lncRNA DLEU2 as a miR-181a sponge regulates SEPP1 protein and inhibited muscle differentiation and regeneration through biological information analysis and vitro experiments. This study would like to provide a new therapeutic target for treating sarcopenia caused by aging through detecting the lncDLEU2/miR-181a/SEPP1 pathway.

## Results

### DE-mRNAs from GEO datasets

After normalization, 11 differentially expressed mRNAs were obtained from skeletal muscle samples (GSE8479, GSE1428 and GSE52699) (Figure 1A). Multiple volcano plots of differential expression are presented in Figure 1B. For those data, SEPP1 were the co-upregulated mRNAs and GFOD1, GOT1, and SV2A were the co-downregulated mRNAs in the sarcopenia group. The expression levels of mRNAs acquired from GSE8479, GSE1428 and GSE52699 datasets are shown in Figure 1C.

**Figure 1.**
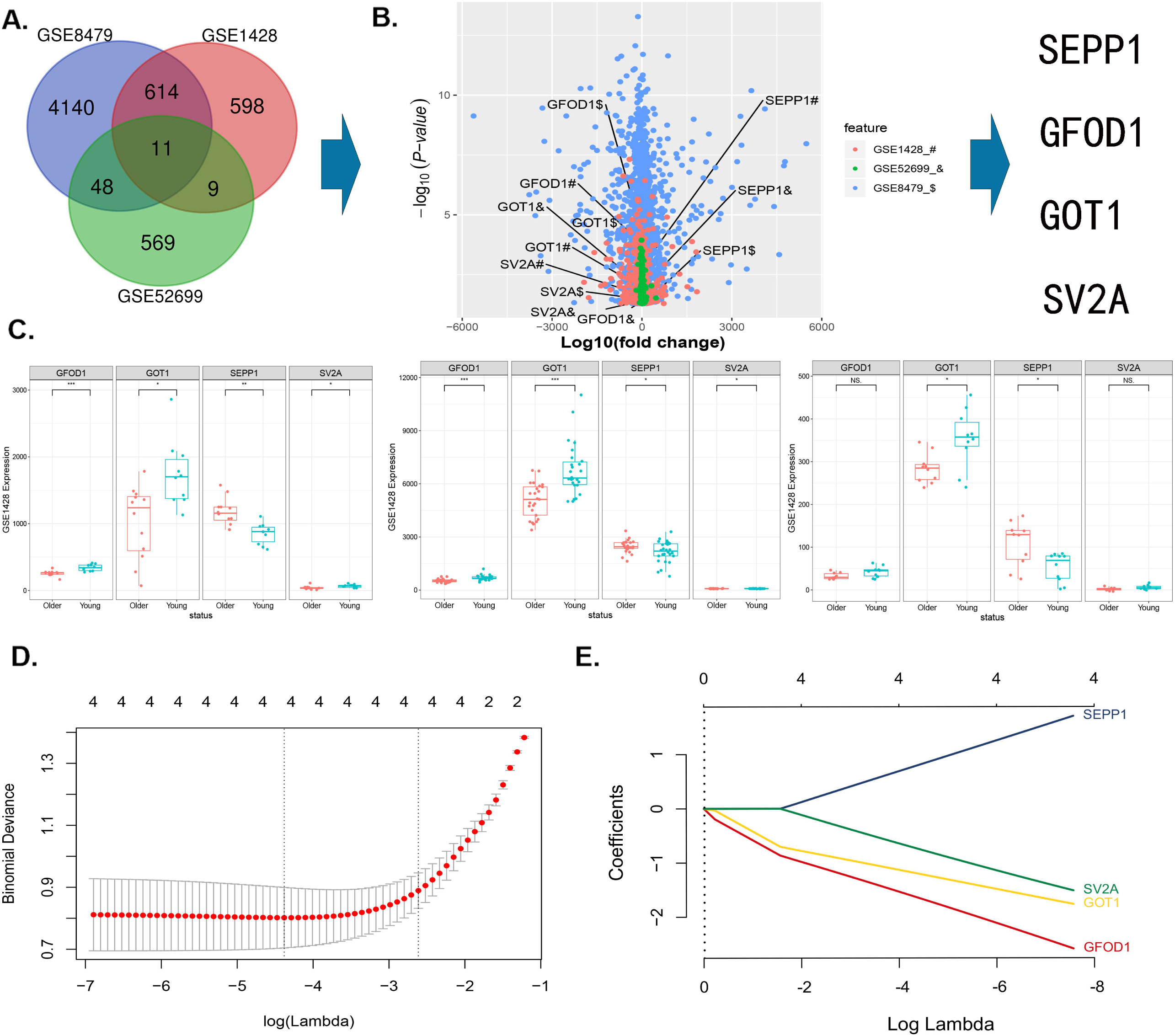
A. Veen diagram demonstrates the intersection of different expression mRNAs datasets of GSE8479, GSE1428, and GSE52699. B. Multi-volcano plot of DE-mRNAs from GSE8479, GSE1428, and GSE52699. C. Association between the expression of DE-mRNAs (SEPP1, SV2A, GOT1 and GFOD1) in different datasets.*P<0.05 D. DE-miRNAs selection by means of the LASSO regression model. The selection of the optimal parameters (lambda) in the LASSO model uses the minimum criterion of 5-fold cross-validation. A dashed line was drawn at the best value using the minimum criterion and 1se (standard error) of the minimum criterion. E. LASSO coefficient profiles of the 4 features. Generate a coefficient outline based on the log (lambda) sequence, where the optimal lambda gets the characteristics of four non-zero coefficients.

### mRNAs selected by predictive modeling of sarcopenia risk

Based on the LASSO regression model, we selected 4 key differentially expressed mRNAs (DE-mRNAs) predictors of sarcopenia risk from 4 features: SEPP1, GFOD1, GOT1, and SV2A (Figure 1D,E). A nomogram model containing these independent predictors is shown in Figures 2A and C. The calibration curve was well matched, and the AUC of training GEO database cohort was high to 0.914 (Figure 2E,F). Besides, Our clinical data demonstrated that the predictive model had an AUC of 0.790 (Figure 2F), suggesting that the nomogram could be used to forecast the chances of sarcopenia. The C index is 0.915(95% CI, 0.84052-0.98948)in the training set, 0.923 (95% CI, 0.86616-0.97984)in the validation set and was validated to be 0.79(95% CI,0.66848-0.91152) by clinic cohort, suggesting strong discriminatory power and accurate predictive performance. The decision curve analysis of the nomogram is shown in Figure 2G. Those implied that the nomogram model can be a tool for clinical practice, including early intervention of risk factors for sarcopenia, thereby reducing the risk of sarcopenia. In total, matrix data for DE-mRNAs and predicted risk score for sarcopenia from the clinic cohort and the entire GEO database cohort were presented as heatmap and risk plot in Figure 2B and Figure 2D.

**Figure 2.**
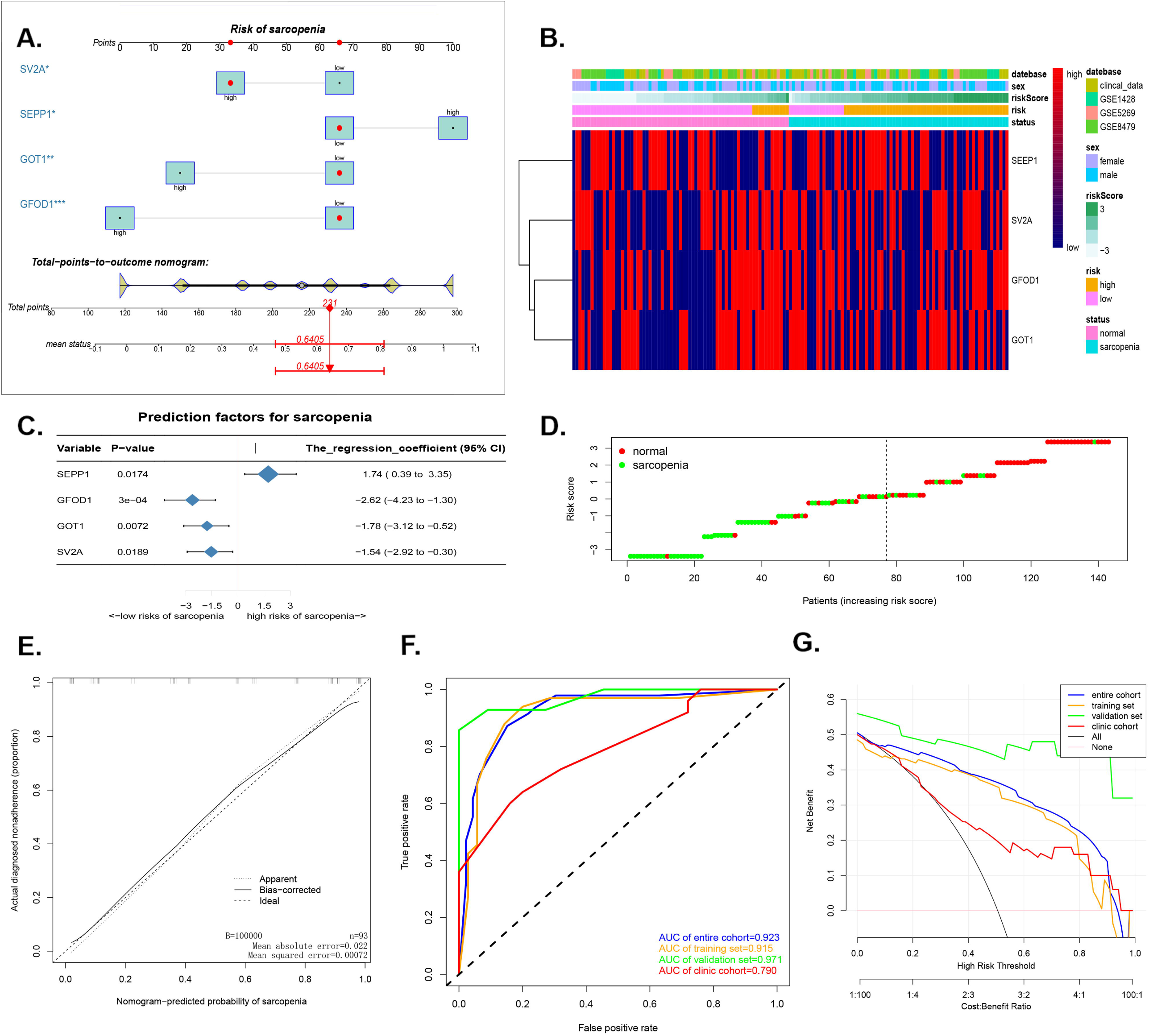
A. The nomogram model of sarcopenia. Note: 4 DE-miRNAs including the expression level of SEPP1, SV2A, GOT1 and GFOD1 were included. B. Heapmap plots of DE-mRNAs (SEPP1, SV2A, GOT1 and GFOD1) in the samples from clinic and datasets of GSE8479, GSE1428, and GSE52699. C. The forest chart of prediction factors for sarcopenia. D. Analysis of DE-mRNAs risk score of sarcopenia in the samples from clinic and datasets of GSE8479, GSE1428, and GSE52699. E. A calibration curve for the predictive model of sarcopenia. Note: The x-axis is the risk of skeletal muscle reduction. The y-axis represents the actual incidence of sarcopenia. Diagonal dashed lines represent perfect predictions for an ideal model. The solid line indicates the prediction ability of the prediction model. The more the solid line matches the dotted line, the better the prediction ability. F. The AUC of the sarcopenia normogram is equal to the accuracy of randomly selected samples. G. Decision curve for intervention benefit probability. The figure shows the decision analysis curve for training set, validation set, the entire cohort and the clinic cohort.

### ceRNA network was established

Differentially expressed lncRNAs (DE-lncRNAs) and differentially expressed miRNAs (DE-miRNAs) were obtained using the parallel method described above.Volcano plots of those data were presented in Figure 3A,B and C. Based on databases of miRcode, miRWalk3.0 and miRTarBase, ceRNA network was established in Figure 3D and E. Two DE-lncRNAs (TTTY9B and BCYRN1) and five DE-miRNAs (miR-222, miR-181a, miR-141, miR-137 and miR-101) were down-regulation in the sarcopenia group. And two DE-lncRNAs (DLEU2 and HULC) and seven differentially expressed miRNAs (miR-98, miR-7, miR-218, miR-215, miR-206, miR-203 and miR-195) were up-regulation in the sarcopenia group. Combined with the nomogram prediction model results, these results indicate that SEPP1, GFOD1, GOT1, and SV2A may be key modulators of sarcopenia.

**Figure 3.**
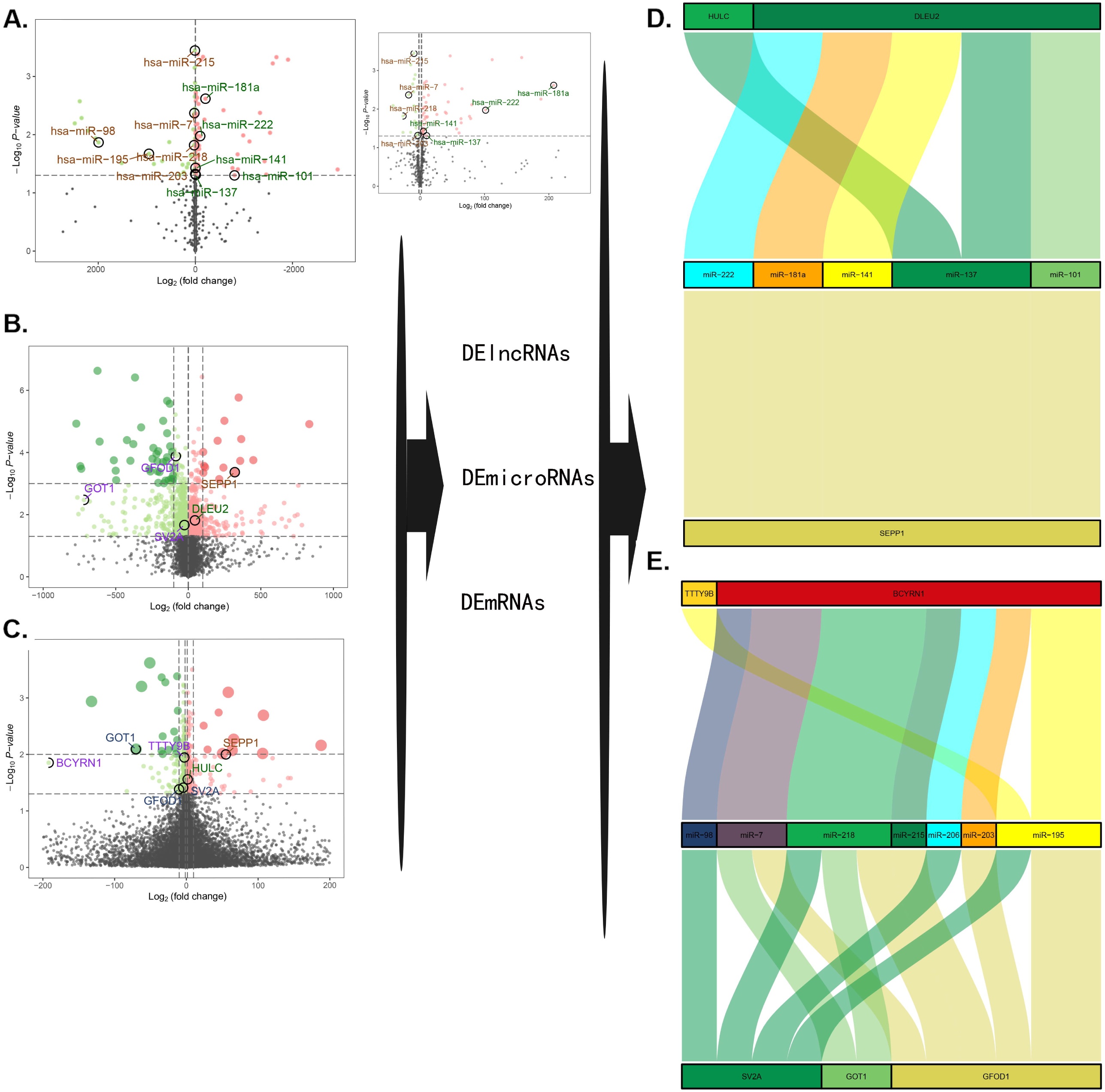
A-C. A showed volcano plot of GSE23527 with 886 miRNAs, and 100 miRNAs were identified either up- or down-regulated in all. B showed volcano plot of GSE1428 with 12427 mRNAs and lncRNAs; 1232 mRNAs and lncRNAs were identified either up- or down-regulated in all. C showed Volcano plot of GSE52699 with 34663 mRNAs and lncRNAs; 637 mRNAs and lncRNAs were identified either up- or down-regulated in all. **Note**: the red dots represent the upregulated ones, the green dots represent the downregulated ones, and the black dots represent the ones that are not significantly differentially expressed in in old muscle samples. D-E. The ceRNA network of sarcopenia.

### Expression and correlation analysis of DE-mRNAs, miR-181a and DLEU2

DE-miRNAs (SEPP1, GFOD1, GOT1, and SV2A),miR-181a and DLEU2 expression levels were examined in the muscle of 25 patients with sarcopenia and 25 patients without sarcopenia by quantitative real-time PCR (Figure 4A). The average SEPP1 and DLEU2 expression level was meaningfully higher in the group of sarcopenia than in the group of control, while the expression of GFOD1, GOT1, SV2A and miR-181a was significantly lower in sarcopenia than the other ones.

**Figure 4.**
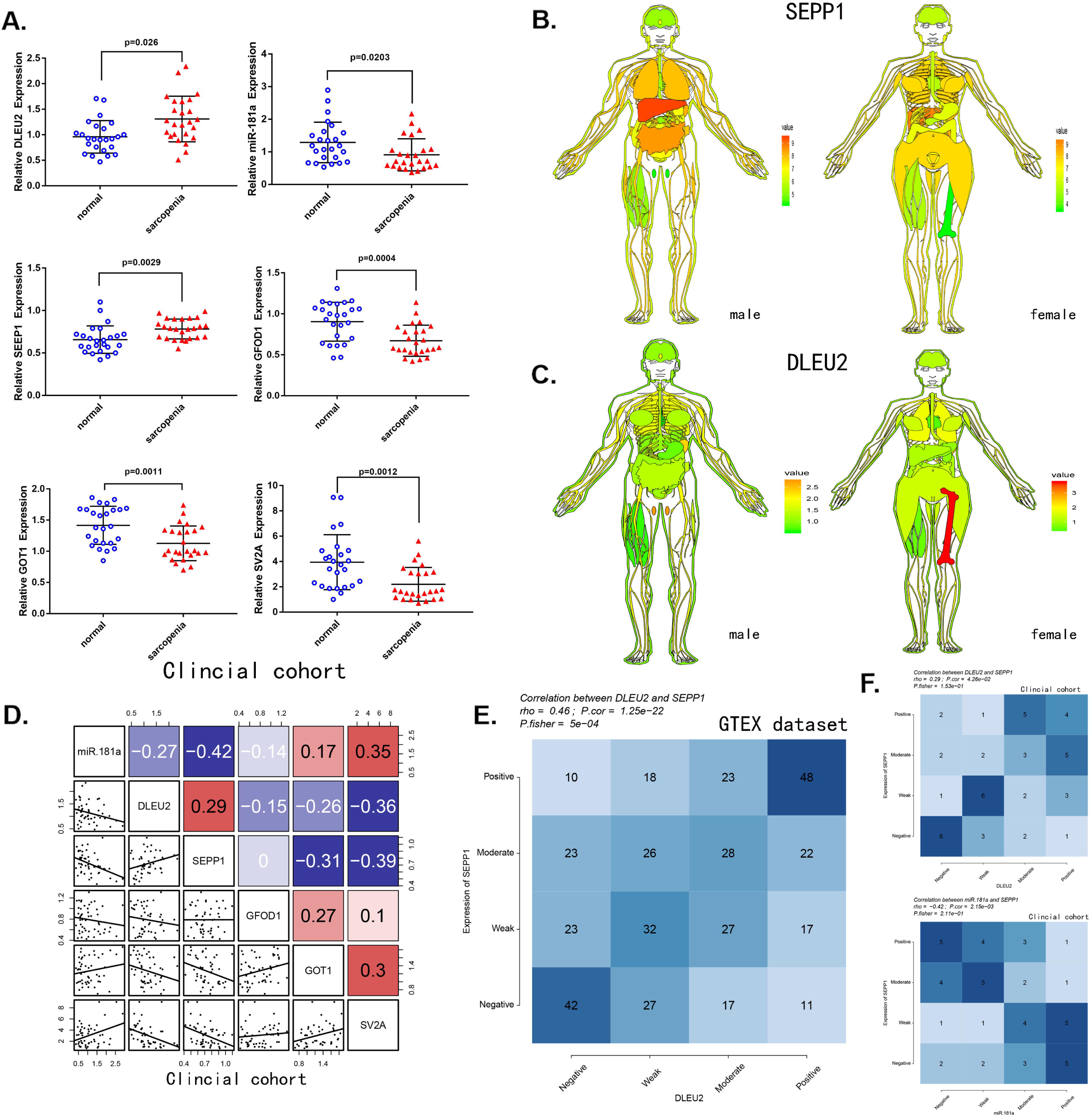
A. Relative expression of DLEU2, miR-181a, SEPP1, SV2A, GOT1 and GFOD1 detected by qPCR in the clinic cohort. B. SEPP1 are lowly expressed in the muscle (GTEX cohort; n =7858). Human tissue-enriched protein expression map of SEPP1 expression. C. DLEU2 are lowly expressed in the muscle (GTEX cohort; n =7858). Human tissue-enriched protein expression map of DLEU2 expression. D. Correlations among DLEU2, miR-181a, SEPP1, SV2A, GOT1 and GFOD1 levels in muscle tissues (clinic cohort; n = 50) E-F. Comparison of the DLEU2 expression scores with the SEPP1 or miR-181a scores in muscle tissues. The correlations are shown in the clinic cohort of muscle tissues (n=50) and GTEX cohort of normal muscle tissues (n=396; c).

The result of ceRNA network (Figure 3D,E) identified that lncRNA DLEU2 acts as a sponge of miR-181a to up-regulation the expression of SEPP1 as well as in the correlation analysis. Firstly, correlation analysis of clinic cohort showed that miR-181a had a significant negative correlation with SEPP1 and DLEU2, combined with (Figure 4D). Besides, SEPP1 also had a significant positive correlation with DLEU2 both in GTEX database and clinic cohort(Figure 4E,F). Furthermore, to understand the expression of SEPP1 and DLEU2 more intuitively, we constructed human tissue–enriched protein expression maps. SEPP1 and DLEU2 were relatively low enriched in muscle (Figure 4B,C). Thus, miR-181a could be a protective factor, whereas DLEU2 and SEPP1 could be detrimental to skeletal development.

### DLEU2 inhibited myogenic proliferation and differentiation of C2C12 myoblasts

To demonstrate the functions of DLEU2, our laboratory group constructed a lentiviral vector encoding with DLEU2 or DLEU2 shRNA, then prepared a lentivirus system for C2C12 cells infection. The results showed that in C2C12 cells transduced by lentivirus DLEU2 (Figure 5A), DLEU2 expression was highly expressed; while shDLEU2-1(shRNA-1) had the highest knockout efficiency in those C2C12 systems (Figure 5C). Besides, the levels of DLEU2 and differentiation markers of myofibrils (MyoD and MyoG) were negative correlation as determined by quantitative RT-PCR analysis (P < 0.05, Figure 5A,C). CCK-8 and EDU assays demonstrated that treatment with DLEU2 reduced cell proliferation and the level of EDU-positive C2C12 cells (Figure. 5B,D and E). Overexpression of DLEU2 in C2C12 cells can significantly reduce the protein and mRNA levels of muscle-derived markers (MyoG and MyoD) as well as SEPP1 and inhibit the proliferation of C2C12 cells, while shDLEU2 (shRNA-1&2) promote these levels in C2C12 cells. (P < 0.05, Figure 5 B,C,F,G and H)

**Figure 5.**
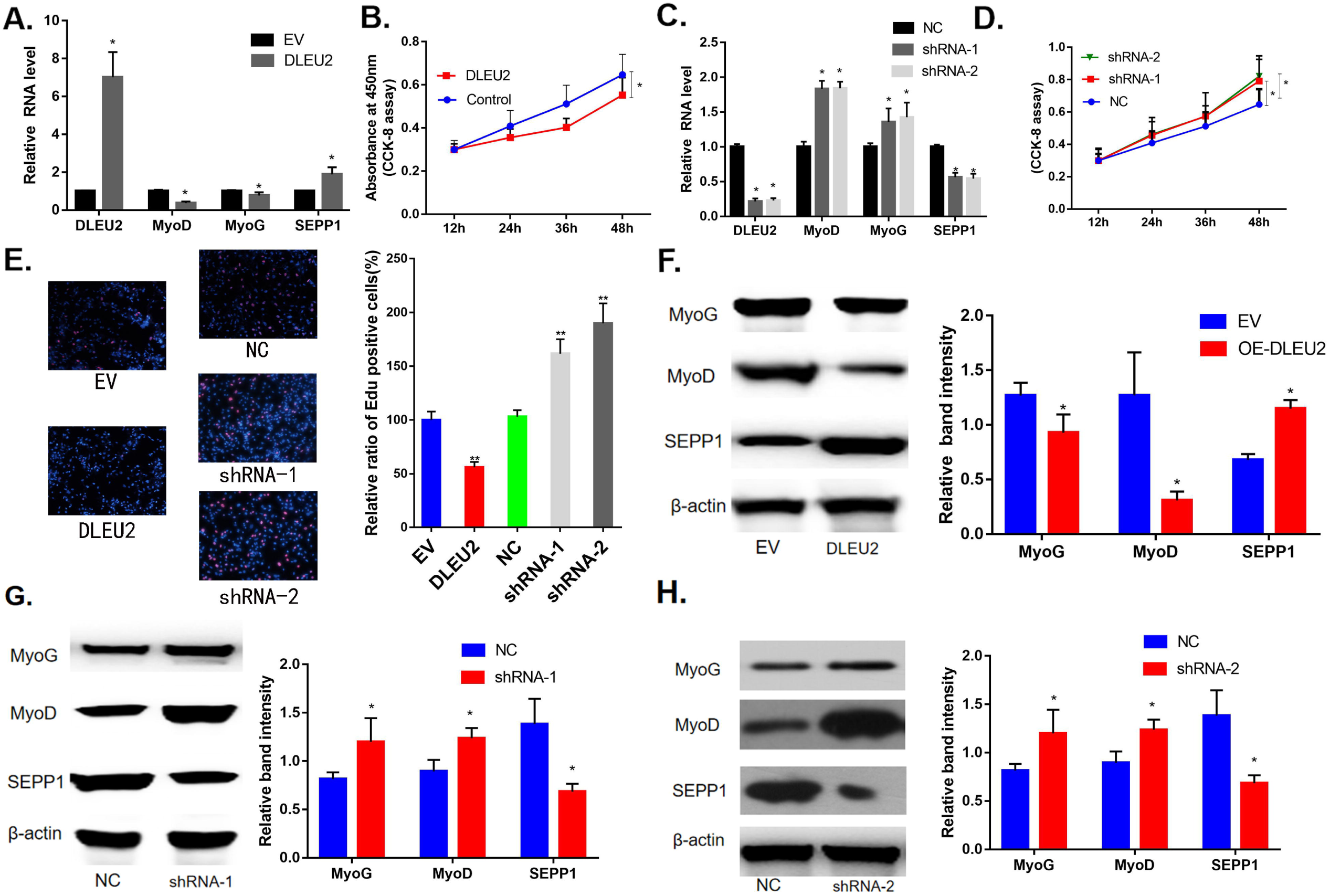
A. SEPP1 and myogenic markers (MyoD and MyoG) mRNA expression levels in DLEU2 infected C2C12 cells by RT-PCR analysis. B. Proliferation of C2C12 cells following the evaluation of the overexpression of DLEU2. C. mRNA expression levels of SEPP1, MyoD and MyoG in DLEU2 shRNA infected C2C12 cells by RT-PCR analysis. D. Proliferation of C2C12 cells following the evaluation in DLEU2 shRNA. E. C2C12 myoblasts were treated with DLEU2 or shRNA-1/2. Cells were stained with Edu. The relative ratio of Edu+ C2C12 cells was quantified. The data represents the mean ± SD (n = 3). Different from control or NC, ** p < 0.01,***p<0.005. F-H. Protein expression levels of SEPP1, MyoD and MyoG in DLEU2 or DLEU2 shRNA infected C2C12 cells by western blot analysis

### Validation of miR-181a targets DLEU2 in C2C12 cells

To more understand the biological mechanism of DLEU2 regulating muscle differentiation, we predict that microRNA181a (miR-181a) is one of the target miRNAs of DLEU2 and a binding site between miR-181a and DLEU2 was also predicted by using RNA hybrid 2.12 (https://bibiserv.cebitec.uni-bielefeld.de/rnahybrid/) (Figure 6A). Further study revealed that the biotinylated miR-181a specifically pulled down the expression level of DLEU2 in C2C12 cells(Figure. 6B). And the double luciferase reporting experiment showed that DLEU2 transfection could reduce the luciferase activity of miR-181a-WT, but did not affect the luciferase activity of miR-181a-Mut. At the same time, over expression of DLEU2 containing the mutant binding site(DLEU2-Mut) did not decrease the luciferase activity of microRNA181a-Mut and microRNA181a-WT (Figure 6C). Co-transfection with microRNA181a-WT upregulation the protein and mRNA levels of muscle-derived markers but decreased the levels of SEPP1 in DLEU2 transfected C2C12 cells (Figure 6D,E). In addition, EDU and CCK-8 assays was also performed to exam the proliferation of C2C12 cells after overexpression of DLEU2 and co-transfection with the microRNA181a mimic or microRNA181a inhibitor. Cells transfected with DLEU2 and treated with the miR-181a inhibitor showed significantly decreased proliferation compared with the others, whereas proliferation was increased in cells treated with the miR-181a mimic (Figure 6F,G). Therefore, SEPP1 protein expression could be increased by DLEU2 infection in C2C12 cells, and miR-181a mimic decreased SEPP1 protein expression as well as promoting the muscle differentiation of C2C12 cells.

**Figure 6.**
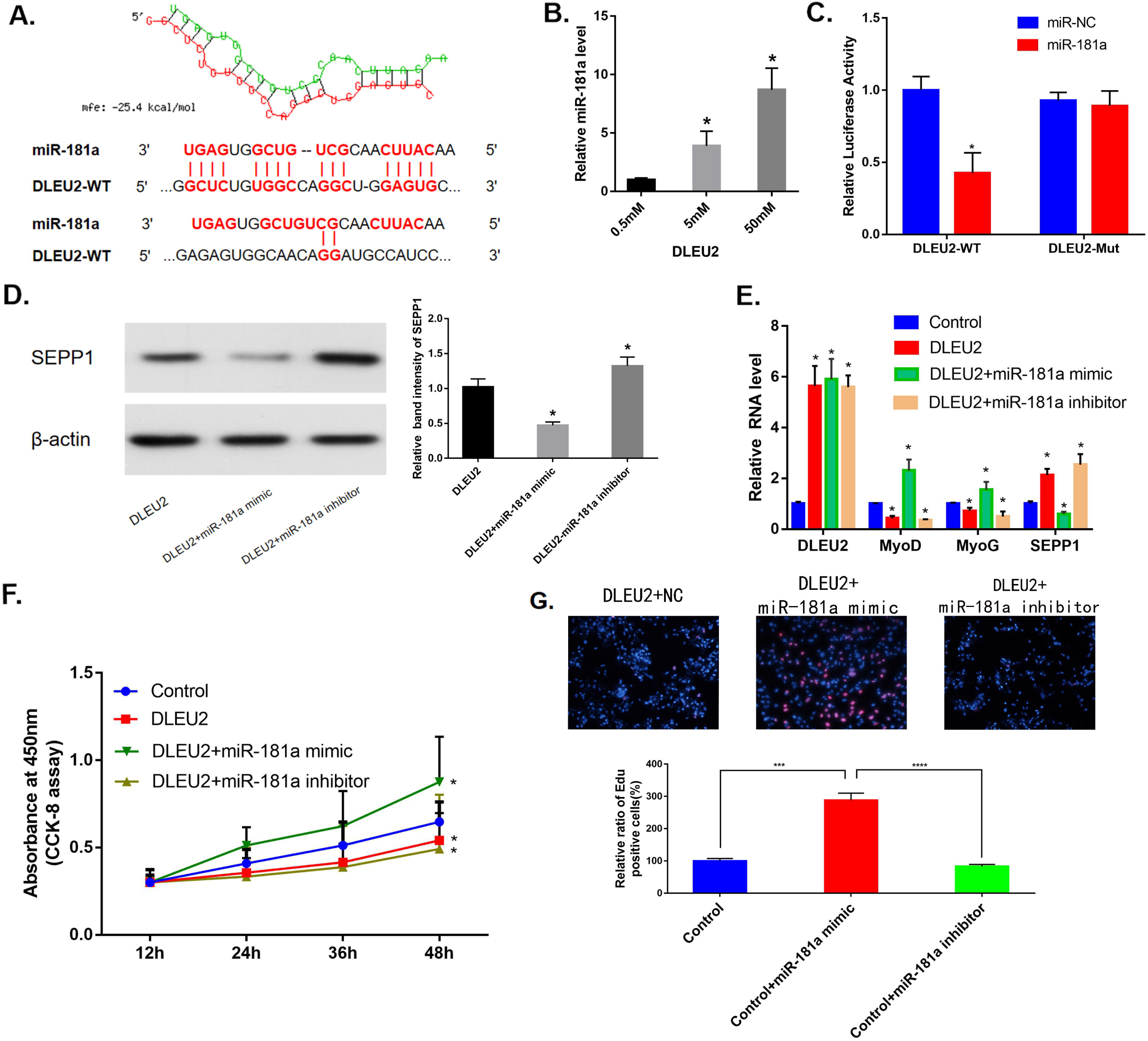
A. RNAhybrid 2.12 predicted the bioinformatics prediction of miR-181a as a target miRNA of DLEU2. MFE: Minimum free energy. B. After transfecting C2C12 cells with different doses of biotin-labeled DLEU2, miR-181a level pull-down experiments and real-time PCR analysis. * P < 0.05 vs. 0.5 mM. C. Luciferase reporter assays to evaluate miR-181a regulation by DLEU2. *P < 0.05 vs. miR-NC. D-E. Real-time PCR and Western blot analysis of mRNA and protein expression of SEPP1, MyoD and MyoG in C2C12 cells co-transfected with mir-181a-WT or mir-181a-Mut with enhanced DLEU2 expression. Data are presented as mean ± SD. U6 small nuclear RNA is used as the internal control of lncRNA and miRNA. GAPDH is used as a messenger RNA control mRNA. F. Proliferation of C2C12 cells after DLEU2 overexpression, miR-181a inhibitor and miR-181a mimic were evaluated. G. C2C12 myoblasts were treated with DLEU2+miR-181mimic or DLEU2+miR-181inhibitor. Cells were stained with Edu. The relative ratio of Edu+ C2C12 cells was quantified. The data shows the mean ± S.D. (n = 3). ***p<0.005,****p<0.0005.

### Characterization of the SEPP1 subtypes regarding different functional pathways

GO function analysis of GSEA showed remarkably effected SEPP1-related signaling functions, such as regulation of skeletal muscle tissue development, skeletal muscle tissue development, skeletal muscle organ development, skeletal muscle fiber development, skeletal muscle cell differentiation, and skeletal muscle tissue regeneration (Figure 7A). And SEPP1-related KEGG pathways were the Endocrine resistance pathway, Glycolysis / Gluconeogenesis pathway, Inositol phosphate metabolism pathway, Oxidative phosphorylation pathway, Purine metabolism pathways and RNA degradation pathways (Figure 7B). These results suggest that SEPP1 play a vital role in the regeneration and development of muscle as well as associated with the pathways of endocrine resistance and cellular metabolism.

**Figure 7.**
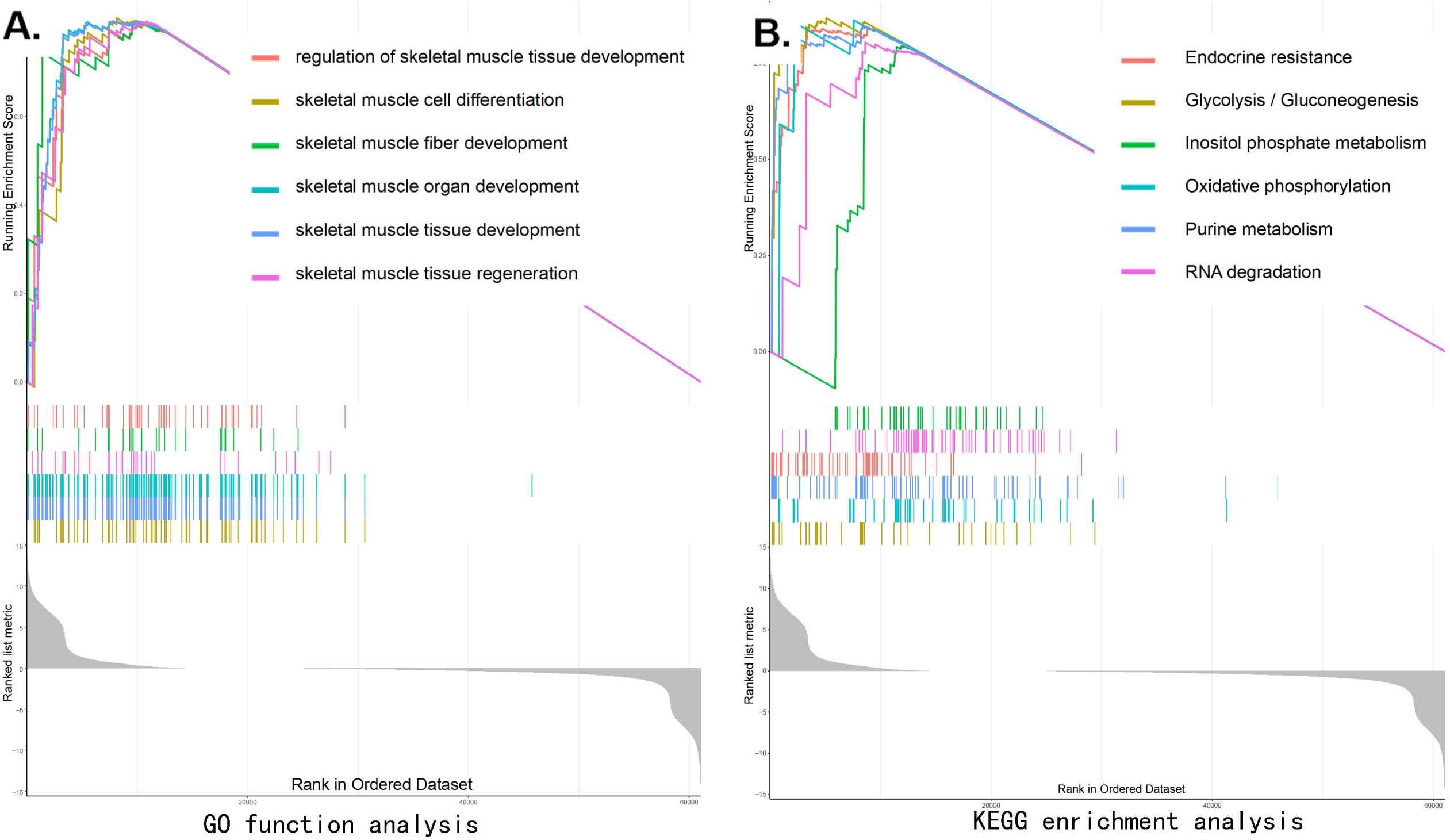
Gene set enrichment analysis revealed the biological pathways and processes related to SEPP1. The enrichment results show that there is a significant correlation between the high and low SEPP1 groups. A. GO enrichment analysis; B. KEGG enrichment analysis

### DLEU2 promoted the expression of SEPP1 protein

MiRWalk and miRcode databases predict that SEPP1 are regulated by miR-181a (Figure 8A). Furthermore, C2C12 cells co-transfected with SEPP1-WT and miR-181a showed less luciferase activity than those co-transfected with SEPP1-WT and microRNA-negative control from our further luciferase reporter assays (NC; p < 0.05) (Figure 8D). In addition, forcing expression of miR-181a in C2C12 cells can meaningfully down-regulate Wnt5a protein expression, while miR-181a inhibitor can up-regulate Wnt5a expression (Figure 8B,C,E). And cells treated with the miR-181a mimic showed significantly promoted proliferation compared with the others (Figure 8F). These results indicated that miR-181a was direct regulators of SEPP1 expression in muscle. In conclusion, lncRNA DLEU2 acts as a miR-181a sponge to regulate the expression of DLEU2, thereby promoting muscle proliferation and differentiation. (Figure 8H)

**Figure 8.**
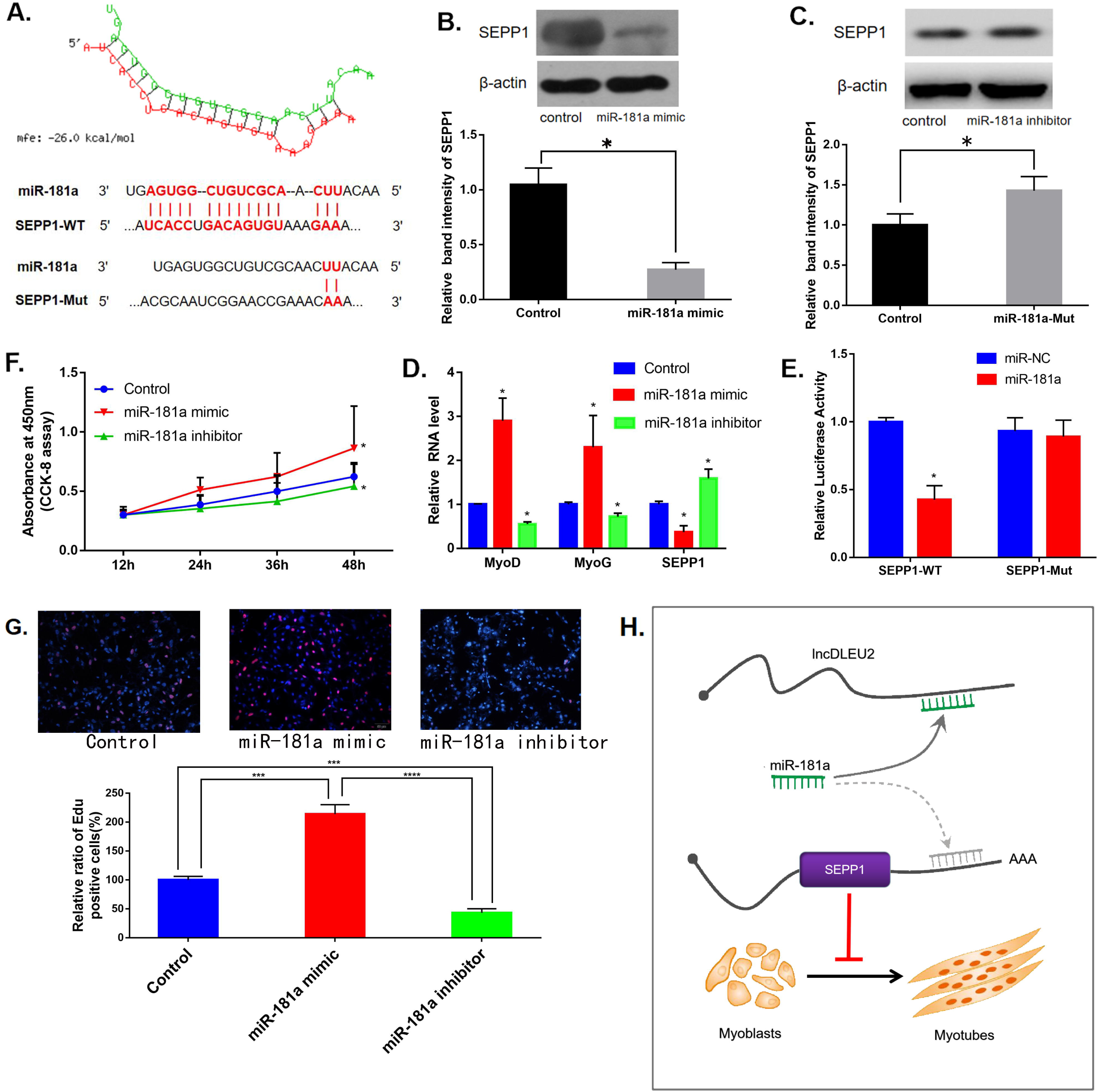
A. Bioinformatic prediction of binding site of miR-181a to DLEU2. B-C. Western blot analysis of SEPP1 in C2C12 cells with miR-181a-Mut or miR-181a-WT. *P < 0.05 vs. control, each trial was assessed in triplicate. D. RT-PCR of SEPP1, MyoD and MyoG in C2C12 cells with miR-181a-Mut or miR-181a-WT. *P < 0.05 vs. control, each trial was assessed in triplicate. E. Luciferase reporter assays to evaluate miR-181a regulation by SEPP1. *P < 0.05 vs. miR-NC. F. Proliferation of C2C12 cells after DLEU2 overexpression, miR-181a inhibitor and miR-181a mimic were evaluated. G. C2C12 myoblasts were treated with miR-181mimic or miR-181inhibitor. Cells were stained with Edu. The relative ratio of Edu+ C2C12 cells was quantified. The data represents the mean ± S.D. (n = 3). ***p<0.005,****p<0.0005. H. **Model of the lncDLEU2-miR-181a-SEPP1 pathway in inhibiting muscle differentiation and proliferation**. Overexpression of lnc-DLEU2, antagonizes miR-181a in competition with SEPP1, to control SEPP1 protein level and inhibiting muscle differentiation and proliferation.

## Discussion

In this research, we found that lncRNA DLEU2 acts as a miR-181a sponge and inhibits skeletal muscle regeneration and differentiation. The lncDLEU2-miR-181a-SEPP1 pathway inhibits muscle differentiation and regeneration could be used as novel therapeutic targets for the treatment of sarcopenia caused by aging. In addition, this study is the first to forecast the chances of sarcopenia based on four molecular markers (SEPP1, SV2A, GOT1 and GFOD1). Besides, the prediction model performed well based on the detection of clinical samples, which can better evaluate clinical efficacy and prognosis[25].

Many studies have shown the important role of the ceRNA network in sarcopenia, and emphasized the need to systematically identify altered mRNAs, miRNAs, and lncRNAs in skeletal muscle development.[16-24] This study first construct a sarcopenia ceRNA network by combining bioinformatics analysis and sarcopenia prediction models. This prediction model uses only four molecular markers, and still has a high C index and AUC value after clinical data validation, which indicates that this tool have the ability to accurately assess the risk of sarcopenia. After that, through the analysis of clinical sample data and the correlation analysis of GTEX database, we selected lncDLEU2-miR-181a-SEPP1 pathway as potential targets for further research. Finally, cell experiments confirmed that DLEU2 as a miR-181a sponge up-regulated SEPP1 inhibited the proliferation and differentiation in C2C12 mouse myotubes. DLEU2 gene is conserved in humans and mice.[26] It is reported that DLEU2 acts as a variety of microRNA sponges (miR-455, miR-496,miR-30c-5p, etc.) and can block cell proliferation in a variety of ways.[27-32]DLEU2 is highly expressed in patients with osteoporosis,[33] while healthy skeletal muscles is necessary for bone prenvent from osteoporosis.[34] Analysis of our clinical samples also suggests that high expression of DLEU2 may be one of the risk factors for sarcopenia in the elderly. Cell experiments show that silencing lncDLEU2 promotes C2C12 differentiation and proliferation, while forcing lncDLEU2 expression is not conducive to the proliferation and differentiation of C2C12 myotubes. These results show that DLEU2 should performance an negative part in skeletal muscle development.

In this study, we predicted that miR-181a contains a DLEU2 binding site through bioinformatics investigation. The luciferase analysis and pull-down method confirmed the direct binding of miR-181a’s response elements to the DLEU2 transcription. Current research shows that miR-181a is important in the establishment of muscle phenotype and significantly expressed during skeletal muscle cell differentiation[35]. MiR-181 inhibits the occurrence of sarcopenia mainly through three ways: (1) miR-181 increases the expression of MyoD and MyoG for myogenic differentiation, and promotes myogenic differentiation and expression of muscle markers[36,37];(2) Increase the proportion of type II muscle fibers in skeletal muscle, thereby increasing skeletal muscle strength[38].Our research also found that lncDLEU2 interacts with miR-181a to reduce MyoD and MyoG after transcription.

The above studies show that microRNA181a plays a protective part in the differentiation and proliferation of skeletal muscle. In our research, a luciferase assays confirmed the binding of microRNA181a as well as SEPP1 in C2C12 cells. Further in vitro studies have also shown that miR-181a may promote muscle cell proliferation and differentiation through targeted down-regulation of SEPP1 protein. As described below, SEPP1 may inhibit muscle cell proliferation and differentiation through multiple pathways. First, SEPP1 inhibit oxidative phosphorylation energy metabolism activation key mediator[39], this may be one of the pathogenesis of sarcopenia. SEPP1 may be detrimental to the growth and development of muscle cells, so studies have found that the expression of SEPP1 in muscle is relatively low compared to other tissues, such as the brain and testes, which is consistent with our results.[40] Second, researchers suggest high expression of SEPP1 may lead to selenium deficiency. [41] Selenium deficiency promotes the body to induce down-regulation of Sepp1 (P <0.05), thereby avoiding muscular dystrophy due to selenium deficiency.[41]

Numerous studies suggest that diabetes is an vital adverse factor for sarcopenia.[42-44] Interestingly, abnormal glucose metabolism in diabetic patients also leads to abnormal upregulation of Sepp1 expression[45], and high expression of SEPP1 may be one of the mechanisms of diabetes causing sarcopenia.

In addition, it has also been reported that SEPP1 is positively correlated with TNF-α levels, and high expression of TNF-α is characteristic of primary muscle disease and high glucose microenvironment.[46,47] In summary, when DLEU2 is knocked out or over-expressed in C2C12 cells, the expression level of miR-181a is up-regulated or down-regulated, resulting in a decrease or increase in the level of SEPP1 protein, and ultimately up or down regulation of muscle proliferation and differentiation, respectively. These data suggest that DLEU2 may interact with miR-181a to up-regulate the level of SEPP1 protein after transcription.

With the increase of age, the muscle mass of the human body gradually decreases. The incidence of sarcopenia is 13% in the elderly aged 60-70, and it reaches 50% at the age of 80[48-50]. Accurate risk assessment will allow doctors to better assess the risk of patients’ illness and facilitate communication between doctors and patients, while also preventing costly medical treatment from occurring. Therefore, we first developed an effective prediction model for sarcopenia, which can provide further theoretical guidance for the treatment and research of sarcopenia; and we also first established a ceRNA network based on the clinical prediction model and verified it experimentally, which has stronger clinical utility.

However, current research also has some limitations. First, due to the limitations of the current data sources, the prediction model will include more factors and samples for further optimization in the future; second, only the most significantly DE-lncRNAs, DE-miRNAs and DE-mRNAs were included in the analysis and ceRNA network construction; third, verification of the ceRNA network is awaiting further animal experiments.

In summary, this study found that lncRNA lncDLEU2 acts as a miR-181a sponge to regulate the SEPP1 protein, thereby inhibiting muscle proliferation and differentiation, which may be a new therapeutic target for reversing aging skeletal muscle atrophy. And based on the four molecular markers (SEPP1, SV2A, GOT1 and GFOD1), a new prediction model with well accuracy can be developed to help clinicians predict the risk of sarcopenia.

## Materials and methods

### Study participants

We identified patients residing in Shanghai, China, who underwent patellar surgery at the Shanghai First People’s Hospital between January 2013 and October 2018. Data for all the patients were collected by the same researcher through appointment-based telephonic interview, outpatient services, and community follow-up. Written informed consent was sought from all enrolled subjects. In this study, we included only fracture patients aged ⩾ 55 years, since postoperative sarcopenia after fractures is rare in young patients. The laboratory muscle tissue test results taken into consideration were those obtained from fine needle aspiration of quadriceps femoris muscle. The study complied with the Declaration of Helsinki and was approved by Shanghai First People’s Hospital (no. 2019SQ059); all subjects provided written informed consent.

The inclusion criteria were as follows: (1) availability of complete data for baseline clinical characteristics (age, body mass index, etc.) and follow-ups, (2) basic communication skills, (3) age ⩾ 55 years, (4) diagnosis of sarcopenia according to the AWGS criteria, (5) patients had been managed with self-care before surgery.

Seventy-five patients were screened, of which 50 were found to be eligible according to the inclusion criteria and availability of the completed questionnaire and the laboratory muscle tissue test. The patients included in the final analysis included 24 males (age: 55–88 years, mean age: 70.3 ± 9.1years) and 26 females (age: 59–86 years, average age: 68.3 ± 7.3 years).

### Methods of assessment

The diagnostic criteria established by the 2014 Asian Working Group for Sarcopenia (AWGS) and EWGSOP2 defines sarcopenia.[51, 52] We used a fixed distance of 6 m, as recommended by AWGS, to measure the subject’s daily walking speed. [53, 54] For patients with a walking speed of ⩽0.8 m/s, we used the bioelectrical impedance analysis (BIA) to assess muscle mass by using a bioimpedance meter (TANITA RD-953, Japan). The results of BIA are very similar to those of double-energy X-ray absorptiometry and magnetic resonance imaging; BIA also offers the advantages of safety, technical simplicity, low cost, and high patient compliance.[55, 56] All the results of BIA have been standardized by using cross-validated Sergi.

### Data retrieval

The dataset supporting the conclusions of this article is available in the GEO database (http://www.ncbi.nlm.nih.gov/geo/). Initially, datasets in which mRNA or miRNA expression were compared between muscle from humans with sarcopenia were included. The abstracts and titles of studies identified were scrutinized, and the full texts of studies that met the criteria were read and evaluated. The GSE23527 miRNA expression array dataset, based on the GPL10358 platform (LC_MRA-1001_miRHuman_11.0_080411 (miRNA ID version)), was selected for further study due to high data quality and relatively large sample size. Those datasets, which contains microarray data from human muscle samples exhibiting sarcopenia and normal and are based on three datasets (GSE8479-GPL2700, Sentrix HumanRef-8 Expression BeadChip; GSE1428-GPL96,[HG-U133A] Affymetrix Human Genome U133A Array; and GSE52699-GPL10558,Illumina HumanHT-12 V4.0 expression beadchip) were also selected. All original files and platform probe annotation information files were saved.

### Identification of differentially expressed genes

All data were normalized using the “normalize between array” function of the “LIMMA” R package from the bioconductor project.[57] This package was also used to identify diferentially expressed lncRNAs (DElncRNAs), mRNAs (DEmRNAs), and miRNAs (DEmiRNAs) between sarcopenia and normal samples from the GSE23527, GSE8479, GSE1428 and GSE52699 datasets.[58] Thresholds set at P < 0.05 and |logFC| > 1.

### Logistic regression of sarcopenia data

The series matrixs from GEO datasets (GSE8479, GSE1428 and GSE52699) were downloaded. Data from skeletal muscles in the older (N = 47) and younger (N = 46) groups were analyzed using R software (version 3.5.3). The samples were randomly divided into training and validation (7: 3) groups. These analyses were put into practice by using the “caret” package to identify and evaluate models. Besides, Our clinical data (25 older, 25 younger) were used to confirm the resulting model. All results were saved in text format for subsequent hierarchical clustering analysis using the Complex Heatmap package.

To identify relevant risk factors in sarcopenia, LASSO method was applied and selecting features with nonzero coefficients. This method is used widely to reduce high-dimensional data.[59-61] A predictive model including the selected features was established using multivariable logistic regression (two-sided P < 0.05).[62] Odds ratios with 95% confidence intervals (CIs) were calculated.

A predictive model was also established for predicting sarcopenia risk based on all potential predictors.[63, 64] For determination of the sarcopenia nomogram, we established calibration curves. Calibration accuracy was assessed statistically using the rms package, with high significance indicating that the model could not provide accurate calibration.[65] Harrell ‘s C-index was calculated to assess the biased performance of the sarcopenia nomogram and corrected by bootstrapping (1,000,000 bootstrap resampling).[65] The clinical utility of the nomogram was assessed by decision curve analysis.[66] After that, the net benefit was calculated as previous study.[67]

### Constructing the ceRNA network

A visual co-expression ceRNA network of DE-lncRNAs, DE-miRNAs, and DE-mRNAs was constructed by using ggalluvial R software package (version :0.9.1).[68] Using the miRcode database, we confrmed the interactions between DElncRNAs and DEmiRNAs.[69] Next, the miRWalk3.0 database(http://mirwalk.umm.uni-heidelberg.de/), which includes 10 databases (Targetscan, RNA22, PITA, PICTAR5, PICTAR4, RNAhybrid, miRWalk, miRDB, miRanda, and DIANAmT), and the miRTarBase(Version 7.0), which comprises validated miRNA target interactions from experiments, were used to assess correlations between DE-mRNAs and DE-miRNAs.[70]

### Analysis of data from the GTEX databases

To clarify the correlation between DElncRNAs and DEmRNAs in ceRNA. The R software (https://www.r-project.org/) with several publicly available packages was used for statistical analysis of data from the GTEX databases. A human tissue – enriched protein expression map and a boxplot of genes were generated using the “gganatogram” and “ggpubr” models, respectively. For the genotypic correlation analysis, the Fisher’s exact test or χ^2^ test (two-sided) was used.

### Quantitative real-time PCR (qPCR)

Total mRNA and lncRNA was isolated from cell cultures using the Mini-BEST Universal RNA Extraction kit (TaKaRa, Kyoto, Japan), following cDNA synthesis using the Prime-Script RT Master Mix (TaKaRa). Finally, the qPCR assays were detected using the SYBR Green Master Mix (TaKaRa) with PCR LightCycler480 (Roche Diagnostics, Basel, Switzerland).

Total miRNA was isolated from cell cultures using TRIzol® reagent (Gibco/Life 270 Technologies, Thermo Fisher Scientific). miRNA quantity and quality were detected by stem-loop quantitative RT-PCR (TaqMan probe method). Purified miRNA was used for first-strand cDNA synthesis with M-MLV reverse transcriptase and primers base on the instructions (Promega, Fitchberg, MA, USA). The primer sequences were designed by Primer Premier and the sequences were as follows: microRNA 181a forward 5’-TGAACATTCAACGCTGTCG-3’ and reverse 5’-GCAGGGTCCGAGGTATTC-3’.

### Western blot analysis

To evaluate protein expression, cells were harvested in RIPA buffer containing a protease inhibitor cocktail, and total protein was quantified using a bicinchoninic acid kit (Pierce, Rockford, IL, USA). Aliquots were separated and then electroblotted onto a 0.45-µm PV membrane (Immobilon™; Merck Millipore, Darmstadt, Germany). The membranes were blocked and then probed overnight with the primary antibodies anti-SEPP1 (1:1000, #ab193193; Abcam, USA), anti-MyoD (1:1500;Invitrogen, Carlsbad,CA,USA), anti-MyoG(1:1500;Invitrogen,Carlsbad,CA,USA), anti – β -catenin (1:5000, #ab32572; Abcam, USA), and anti – active β -catenin (1:500, #05-665; Merck Millipore).

### Cell culture and differentiation

C2C12 cells were obtained from American Type Culture Collection (ATCC, CRL-1772™,Manassas, VA,USA). The cells were cultured at a confluent density in growth medium and consisted of Dulbecco’s modified eagle medium(DME-M), 10% heat-inactivated fetal calf serum (Biowest, St. Louis, MO), and 1% penicillin-streptomycin. C2C12 cells were differentiated into myocytes or myotubes in a differentiation medium, consisting of DMEM containing 2% heat-inactivated horse serum (Invitrogen, Carlsbad, CA,USA) and 1% penicillin-streptomycin. All these cells were maintained in a humidified atmosphere at 37 ° C, 5% CO2.[71]

### Cell Counting Kit-8 (CCK-8) determination

After transfection cultured in growth medium, cell proliferation was observed using TransDect CCK (TransGen Biotech, Beijing, China) according to the protocol. The absorbance was measured with optical density using a Model 680 Microplate reader (Bio-Rad, Hercules, California, USA).

### Cell transfection

According to a previously reported modification protocol, C2C12 cells were cultured to 60% confluence.[73] The culture medium was then removed and 1.5×108 IU virus particles were added with 8 g/mL hexadimethyl bromide (Sigma-Aldrich, St. Louis,MO, USA). After that, DMEM + virus particles with 10% were changed to DMEM fiber channel standard and the cells were cultured for 1-7 days.

### Construction of lentiviral vectors and lentivirus production

To construct the lncDLEU2 overexpression lentiviral vector, we subcloned DLEU2 and the full-length lncDLEU2 into the lentiviral GV112 vector according to the manufacturer’s instructions.[74] This vector was provided by Shanghai Genechem (Shanghai, China). For the lncDLEU2-KD lentiviral vector, the shRNA subcloned of the lncDLEU2 or negative control scramble sequence was used in the GV112 carrier. Shanghai Genechem designed two shRNA sequences (shDLEU2-1: 5’-AGCTCAGATTCTCTCCTTT-3’, shDLEU2-2: 5’-TGAAAGGTGTACTGCAAGGAA-3’). The lentivirus expression vector was co-transfected into C2C12 cells using TransIT-LT1 (Mirus Bio). After transfection, supernatants were collected at 48h and 72h, concentrated by ultracentrifugation at 25,000 rpm for 90 minutes, and recovered in an appropriate volume of OptiMEM (Gibco, Waltham, MA, USA). Determination titer (IU / mL) of infected particles by real-time qPCR.[75]

### Transfection of miRNAs

Transfection of miRNAs was performed as previously described.[76] In short, miR-181a is enhanced and inhibited using chemically synthesized miRNAs mimics and inhibitors (Gene Pharma (Shanghai, China)). According to the manufacturer’s protocol (Ribobio, Guangzhou, China), 24 hours after seeding, cells were transfected using a riboFECT™ CP transfection kit for 24 hours. The transfection efficiency was measured through real-time quantitative PCR after transfection 48 hours.

### Pulling down the Biotin-labeled lncDLEU2

Biotin-labeled lncDLEU2 was synthesized by Sangon Biotech (Shanghai, China).Incubate different doses of biotin-labeled DLEU2 (0.5 mM, 5 mM, and 50 mM) with cytoplasmic lysate of C2C12 cells transfected with miR-181a for 30 minutes at room temperature. And the complexes were isolated by the streptavidin-coated magnetic bloodworm (Dynal, Waltham, MA, USA). The captured RNA is purified and subjected to real-time PCR analysis after completing the washing step.[77]

### Dual luciferase reporter assay

According to the previously report, the putative sequence of the miR-181a binding site and the mutant sequence were cloned into the pmirGlO dual luciferase miRNA target expression vector (Promega, Madison, WI, USA) to form a reporter vector. 41 Co-transfecting the reporter vector with lncDLEU2-Mut or lncDLEU2-WT into C2C12 cells using Lipofectamine 2000 (Invitrogen, Carlsbad, CA, USA), similarly steps when co-transfect the reporter vector with SEPP1-WT or SEPP1-Mut. According to the manufacturer’s protocol, the dual luciferase reporter assay system (Promega, Madison, WI, USA) was used to detect renilla luciferase activities after 48h.

### EDU assays

The C2C12 cells treated under different treatment conditions were seeded in 24-well plates at a rate of 1 × 105 cells / well and incubated for 24h. As previously reported basing on the manufacturer’s instructions, the 5-erhynyl-20-deoxyuridine (EDU) incorporation assay was performed with an EDU assay kit(#COO75S, Beyotime Biotechnology). The proportion of EDU-positive cells was counted according to the images taking under a laser scanning confocal microscope (Olympus)[78-80].

### Gene set enrichment analysis(GSEA)

GSEA is a “molecular signature database” used to investigate potential mechanisms by using the project of JAVA (http://software.broadinstitute.org/gsea/index.jsp). [81] The number of random samples was set to 1000, and the significance threshold was set to p <0.05.

### Statistical analysis

Statistical analysis was using GraphPad Prism (version 7.0) software. Results are expressed as the mean ± standard deviation of three or six independent experiments. Statistical significance used one-way analysis of variance or two-tailed t test. Correlation analysis was performed using pear in correlation test. The difference was statistically significant, *P <0.05.

## Acknowledgements

Thanks for the anonymous reviewers for their valuable comments and suggestions that helped improve the quality of our manuscript.

## Funding

No funding

## Competing interests

The authors declare that they have no competing interests.

## Availability of data and materials

Please contact author for data requests.

## Ethics approval and consent to participate

This study was approved by the Institutional Ethics Review Board of Shanghai General Hospital, Shanghai Jiao Tong University, Shanghai, China.

## Abbreviations

(BMD): bone mineral density
(BPs): biological processes
(CCs): cellular components
(ceRNA): competing endogenous RNA
(CIs): confidence intervals
(DE-miRNAs): differentially expressed micro-RNAs
(DEMs): differentially expressed mRNAs
(EDU assays): 5-ethynyl-2’ -deoxyuridine assays
(EV): empty vector
(GEO): Gene Expression Omnibus
(GFOD1): glucose-fructose oxidoreductase domain containing 1 protein
(GTEX): Genotype-Tissue Expression
(GO): gene ontology
(GSEA): Gene Set Enrichment Analysis
(LASSO): the least absolute shrinkage and selection operator
(lncRNAs): long noncoding RNAs
(NC): normal control
(KEGG): Kyoto Encyclopedia of Genes and Genomes
(mRNAs): expressed messenger RNAs
(PPI): protein – protein interaction
(SEPP1): SELENOP protein
(SV2A): synaptic vesicle membrane protein 2A

## Notes

### Competing Interest Statement

The authors have declared no competing interest.

